# Transcriptomic responses associated with carbon and energy flows under high salinity stress suggest the overflow of acetyl-CoA from glycolysis and NADPH co-factor induces high lipid accumulation and halotolerance in *Chlorella* sp. HS2

**DOI:** 10.1101/817551

**Authors:** Jin-Ho Yun, Michaël Pierrelée, Dae-Hyun Cho, Urim Kim, Jina Heo, Dong-Yun Choi, Yong Jae Lee, Bongsoo Lee, HyeRan Kim, Bianca Habermann, Yong Keun Chang, Hee-Sik Kim

**Affiliations:** Cell Factory Research Center, KRIBB, Yuseong-gu, Daejeon, 34141, Republic of Korea; Aix-Marseille University, CNRS, IBDM UMR 7288, 13009 Marseille, France; Department of Environmental Biotechnology, UST, Yuseong-gu, Daejeon, 34113, Republic of Korea; Department of Microbial and Nano Materials, College of Science and Technology, Mokwon University, Seo-gu, Daejeon, 35349, Republic of Korea; Plant Systems Engineering Research Center, KRIBB, Yuseong-gu, Daejeon, 34141, Republic of Korea; Advanced Biomass R&D Center, 291 Daehak-ro, Yuseong-gu, Daejeon, 34141, Republic of Korea; Department of Chemical and Biomolecular Engineering, KAIST, Yuseong-gu, Daejeon, 34141, Republic of Korea

**Keywords:** acetyl-CoA, *Chlorella* sp. HS2, halotolerance, lipid synthesis, photosynthesis, RNA-seq

## Abstract

Previously, we isolated *Chlorella* sp. HS2 (referred hereupon HS2) from a local tidal rock pool and demonstrated its halotolerance and relatively high biomass productivity under different salinity conditions. To further understand acclimation responses of this alga against high salinity stress, we performed transcriptome analysis of triplicated culture samples grown in freshwater and marine conditions at both exponential and stationary growth phases. *De novo* assembly followed by differential expression analysis identified 5907 and 6783 differentially expressed genes (DEGs) respectively at exponential and stationary phases from a total of 52770 transcripts, and the functional enrichment of DEGs with KEGG database resulted in 1445 KEGG Orthology (KO) groups with a defined differential expression. Specifically, the transcripts involved in photosynthesis, TCA and Calvin cycles were downregulated, whereas the upregulation of DNA repair mechanisms and an ABCB subfamily of eukaryotic type ABC transporter was observed at high salinity condition. In addition, while key enzymes associated with glycolysis pathway and triacylglycerol (TAG) synthesis were determined to be upregulated from early growth phase, salinity stress seemed to reduce the carbohydrate content of harvested biomass from 45.6 dw% to 14.7 dw% and nearly triple the total lipid content from 26.0 dw% to 62.0 dw%. These results suggest that the reallocation of storage carbon toward lipids played a significant role in conferring the viability of this alga under high salinity stress, most notably by remediating high level of cellular stress partially caused by ROS generated in oxygen-evolving thylakoids.

**Summary Statement:** Redirection of storage carbon towards the synthesis of lipids played a critical role in conferring the halotolerance of a *Chlorella* isolate by remediating excess oxidative stress experienced in photosystems.

## 1. Introduction

Microalgae exhibit a greater biomass yield than most terrestrial crops and can be grown with excess nutrients in wastewater sources, prompting its industrial utilization as a biofeedstock for the production of nutraceuticals, pharmaceuticals, cosmetics, and biofuels (Hu et al., 2008; Quinn & Davis, 2015; Smith, Sturm, Denoyelles, & Billings, 2010; Unkefer et al., 2017; Yun, Cho, Lee, Heo, et al., 2018). However, commercial production of algal biomass is not yet considered to be economically competitive because of high energy inputs associated with biomass harvesting and downstream extraction of desirable biomolecules (Laurens et al., 2017; Stephens et al., 2010; Valizadeh Derakhshan, Nasernejad, Abbaspour-Aghdam, & Hamidi, 2015). Importantly, the productivity and operational stability of algal cultivation platforms are prone to be compromised by unpredictable meteorological conditions and culture contamination (McBride et al., 2014; Wang et al., 2016; Yun et al., 2019; Yun, Cho, Lee, Kim, & Chang, 2018; Yun, Smith, La, & Keun Chang, 2016), which has led to multifactorial efforts to develop robust algal “crops” under changing environments, just as in the case of conventional agriculture.

Of environmental conditions that determine the productivity of biomass and desirable biomolecules from industrial crops, salinity appears on the top of the list because high crop sensitivity to presence of high concentrations of salts in the soil or waters (Flowers, Troke, & Yeo, 1977; Peng et al., 2014; Yuge Zhang & Liang, 2006). In particular, the extensive application of chemical fertilizer facilitates accumulation of salts in agricultural fields, which in turn could lead to a positive feedback loop by necessitating an increased application of synthetic fertilizer (Yuge Zhang & Liang, 2006). Notably, industrial algal cultivation platforms require continuous provision of nutrient salts with some studies demonstrating the utilization of saline wastewater sources enriched with nitrogenous and phosphorus nutrients as growth media to drive down the costs of commercial operation of algal cultivation systems (Yun, Cho, Lee, Heo, et al., 2018; Yun, Smith, & Pate, 2015; Zhu et al., 2013). In addition, the direct application of salinity stress for algal cultivation systems has been demonstrated as an effective abiotic inducer of high lipid accumulation and an environmental barrier inhibiting the proliferation of undesirable alien invaders in cultivation systems (Church et al., 2017; Kakarla et al., 2018; Lee, Nam, Yang, Han, & Chang, 2016). For instance, Kakarla *et al.* supplemented 60 g/L of NaCl into concentrated *Chlorella* cultures for 48 hr and reported ca. 58% increase in algal lipid productivity, supporting the possibility of deploying high salinity stress as a promising post-treatment for the cultivation systems targeting to produce algal lipids (Kakarla et al., 2018). Moreover, while high salinity stress could act as an effective method of crop protection in reducing freshwater cyanobacterial or ciliate contaminants, it was successfully demonstrated to facilitate algal harvesting by enlarging cellular diameter and increasing algal settling rates (Church et al., 2017; Lee et al., 2016; von Alvensleben, Stookey, Magnusson, & Heimann, 2013). Even though osmosensitivity of algal crops has been acknowledged (Flowers et al., 1977), there is thus a great industrial incentive to exploit algal diversity and especially high tolerance of some algal species to highly saline environment (Yun et al., 2015).

With the apparent advantages of incorporating high salinity stress into the management of industrial algal cultivation platforms, bioprospecting halotolerant algal strains that exhibit high and reliable production of biomass and/or desirable biomolecules was the motivation of our previous study in which a halotolerant *Chlorella* sp. was isolated from a tidal rock pool (Yun et al., 2019). While the remarkable toughness of *Chlorella* under different physical and chemical stress and its recognition as one of a handful of successful industrial crops have been well documented (Fogg, 2001; Yun et al., 2019), this isolated *Chlorella* sp. HS2 (referred to hereupon as HS2) exhibited relatively high growth under a wide range of salinity conditions (i.e., 0-7% (w/v) of supplemental NaCl) compared to reference *Chlorella* strains (Yun et al., 2019). Importantly, substantial shifts in the composition of fatty acid methyl ester (FAME) and the amount of carotenoid pigments under different salinity conditions led us to speculate that elucidating mechanisms behind relatively short-term (i.e., few days) of algal acclimation to high salinity stress would enable maximizing the industrial potential of HS2 by guiding ongoing efforts in metabolic and process engineering (Oh, Chang, & Lee, 2019; Rathinasabapathi, 2000; Yun et al., 2019).

Herein, we report the transcriptome of HS2 grown in freshwater and marine conditions to accomplish mechanistic understanding of algal acclimation to high salinity stress. Triplicated cultures samples were first obtained at exponential and stationary growth phases in freshwater and marine growth media for *de novo* RNA-seq analysis, and the proximate analysis of harvested biomass was additionally performed along with the measure of photosystem II (PSII) activity. Combined together with the results in our previous study, we were able to elucidate how vital metabolic pathways were shifted under high salinity stress, and an important role of redirecting storage carbon towards the synthesis of lipids in conferring the viability of HS2 and remediating high oxidative stress under high salinity stress.

## 2. Materials and Methods

### 2.1. Strain selection and cultivation conditions

*Chlorella* sp. HS2 was previously isolated from a local tidal rock pool, and its high tolerance to a wide range of salinity conditions was acknowledged (Yun et al., 2019). While the results of HS2 cultivation in 1-L cylindrical PBRs were reported in our previous study (Yun et al., 2019), both autotrophic cultures grown in freshwater inorganic medium and in marine inorganic growth medium supplemented with 3% (w/v) sea salt were subjected to transcriptome analysis. These triplicated cultures were grown under pre-determined optimal light and temperature conditions with continuous supplementation of 5% CO_2_ at 0.2 vvm and agitation at 120 rpm.

### 2.2. PSII activity measurement and proximate analysis

While pigment and FAME composition of harvested HS2 biomass in both freshwater and marine conditions were reported previously, photoautotrophically grown cells in exponential and stationary growth phases were subjected to measurements of the photosynthetic parameters *in vivo* using Multi-Color-PAM (Heinz Walz, Germany) (Shin et al., 2017). After adapting cells under dark condition for 20 min, the light response curves of the relative electron transport rate (rETR), the quantum yields of non-photochemical quenching (Y(NPQ)) and nonregulated excess energy dissipation (Y(NO)) were measured in biological triplicates while increasing the actinic light intensities of 440 nm LEDs with a step width of 2 min (Shin et al., 2017). In addition, proximate analysis of the biomass harvested at stationary growth phase was performed in biological triplicates to further elucidate metabolic shifts in HS2 under high salinity stress. Briefly, lipid content of harvested biomass was analyzed by extracting total lipids from freeze-dried biomass with chloroform-methanol (2:1 (v/v)) following a slightly modified version of Bligh and Dyer’s method (Bligh & Dyer, 1959); the protein content was determined using the method of Lowry (Illman, Scragg, & Shales, 2000; Lowry, Rosebrough, Farr, & Randall, 1951); the carbohydrate content was measured using the phenol sulfuric acid method of Dubois et al. (Dubois, Gilles, Hamilton, Rebers, & Smith, 1956; Illman et al., 2000); and the ash content was analyzed gravimetrically after exposing dry biomass to 500 °C in a muffle furnace for 8 hours (Kent, Welladsen, Mangott, & Li, 2015).

### 2.3. RNA extraction, library construction, and Illumina sequencing

Each of salt-stressed and control PBR cultures was harvested during exponential and stationary growth phases by centrifugation at 4500 rpm for 10 min. Total RNA was then extracted using the Trizol reagent (Invitrogen, Carlsbad, CA, USA), according to manufacturer’s instructions. Subsequently, the RNA samples were treated with DNase I for 30 min at 37 °C to remove genomic DNA contamination, and the quantity and integrity of the total RNA were verified using an Agilent 2100 bioanalyzer. The cDNA libraries were developed according to manufacturer’s instructions (Illumina, Inc., San Diego, CA, USA), and sequenced on the Illumina HiSeq 2000 platform at Seeders Co. (Daejeon, Korea) (Liu et al., 2017). In addition, RNA-Seq paired end libraries were prepared using the Illumina TruSeq RNA Sample Preparation Kit v2 (catalog #RS-122-2001, Illumina, San Diego, CA). Starting with total RNA, mRNA was first purified using poly (A) selection or rRNA depletion, then RNA was chemically fragmented and converted into single-stranded cDNA using random hexamer priming; the second strand was generated next to create double-stranded cDNA. Library construction began with generation of blunt-end cDNA fragments from ds-cDNA. Thereafter, A-base was added to the blunt-end in order to make them ready for ligation of sequencing adapters. After the size selection of ligates, the ligated cDNA fragments which contained adapter sequences were enhanced via PCR using adapter specific primers. The library was quantified with KAPA library quantification kit (Kapa biosystems KK4854) following the manufacturer’s instructions. Each library was loaded on Illumina Hiseq2000 platform, and the desired average sequencing depth was met while performing high-throughput sequencing.

### 2.4. *De novo* assembly and analysis

While raw sequencing data were composed of 100 bp paired-end reads, *de novo* assembly was performed using Trinity 2.8.5 (Grabherr et al., 2011). Assembly quality assessment was carried out with BUSCO 3.0.2 (Simão, Waterhouse, Ioannidis, Kriventseva, & Zdobnov, 2015), for which the chlorophyte database of OrthoDB 10 was employed as datasets at an e-value cutoff of 1e-5 (Kriventseva et al., 2018), and high-quality reads were mapped onto genome sequences by using Bowtie2 2.3.5. Thereafter, the quantification of the number of reads (i.e., counts mapped per transcripts) was performed following alignment and abundance estimation of each Trinity script using RSEM 1.3.2 and Bowtie 1.2.2, respectively (Langmead, Trapnell, Pop, & Salzberg, 2009; B. Li & Dewey, 2011). While transcripts with no count across all sampling points were removed, the matrix of counts for unigenes was used for downstream analyses.

### 2.5. DEG analysis and functional annotation

Prior to functional annotation, differential expression analysis (DEA) was performed first to avoid determining the most relevant transcript for each unigene based on unnecessary assumptions at the early stage. In addition, given that quantitative asymmetry between up- and downregulated unigenes was strong, SVCD 0.1.0, which does not assume the lack-of-variation between up- and downregulated unigene counts (Evans, Hardin, & Stoebel, 2017; Roca, Gomes, Amorim, & Scott-Fordsmand, 2017), was used in normalization for unigenes with the mean of raw counts greater than the first quartile (i.e., 5.9 raw counts) as recommended (Roca et al., 2017). To determine DEGs, we used DESeq2 1.20.0 to compute log-fold change (LFC) and adjusted p-values, and the DEGs between exponential and stationary growth phases were especially based on the adjusted p-values for the interaction between medium type and growth phase.

While unigenes with an adjusted p-value lower than 0.01 was considered DEGs, functional annotation of DEGs was performed using Swiss-Prot, Pfam, and Kyoto Encyclopedia of Genes and Genomes (KEGG) databases. First, following Trinotate 3.2.0’s recommendation, we predicted transcript coding regions that could be assigned to putative proteins using TransDecoder 5.5.0 (Haas et al., 2013). Thereafter, homologies were identified using in parallel BlastP from BLAST+ 2.9.0 and *hmmscan* program from HMMER 3.2.1 to get Pfam domains (Camacho et al., 2009; Eddy, 2011). A second BlastP and *hmmscan* were then run from the predicted proteins, and SignalP 5.0b (http://www.cbs.dtu.dk/services/SignalP/) was used to determined eukaryotic signal peptides within transcripts following homologous sequence search on BlastX. While all Blast run was performed against Swiss-Prot database through DIAMOND 0.8.36 (Buchfink, Xie, & Huson, 2015), an e-value cutoff of 1e-10 was used. Then, KEGG cross-references associated with BlastX or BlastP hits were retrieved to assign each Blast hit with KEGG Orthology (KO). Transcripts without a BlastX or BlastP hit were excluded, and a pair of transcript and coding region was removed when the KO of each of corresponding transcript and coding region was not identical. In addition, when one gene had multiple KOs, the mean of average e-values was computed and the KO with the lowest mean was selected as the most relevant KO. While metabolic pathway maps were constructed using KEGG mapper based on the organism-specific search results of *Chlorella variabilis* (cvr) and biological objects for each KO were determined using KEGG BRITE, enrichment was performed by implementing GSEAPreranked from Gene Set Enrichment Analysis with the conda package GSEApy 0.9.15 (Mootha et al., 2003; Subramanian et al., 2005). A term was considered to be significantly enriched when its false discovery rate (FDR) was lower than 0.25.

## 3. Results

### 3.1. Phenotypic shifts of HS2 under high salinity stress

Shifts in growth, FAME and pigment composition of *Chlorella* sp. HS2 during autotrophy in freshwater (i.e., 0% (w/w) of supplemental sea salt) and marine (i.e., 3% (w/w) of supplemental sea salt) media were reported in the previous study (Yun et al., 2019). Briefly, the results indicated a nearly 10-fold decrease in the maximum cell density of the autotrophic PBRs in marine medium at stationary growth phase, whereas only a two-fold decrease in the average dry cell weight (DCW) was observed (Yun et al., 2019) (Supplementary Figure 1). As microscopic observation revealed, a non-proportional decrease in DCW of HS2 under high salinity stress corresponded to roughly 50% increase in cellular diameter or 3.4-fold increase in cellular volume. While previous study also reported substantial decreases in the amount of algal pigments and relatively amount of polyunsaturated fatty acids under high salinity stress (Yun et al., 2019), TEM images of harvested algal cell suggested the formation of large lipid droplets under high salinity stress (Fig. 1): indeed, proximate analysis of harvested biomass indicated a significant increase in lipid content from 25.0 dw% to 62.0 dw% under high salinity stress, contrasting a nearly three-fold decrease in the amount of carbohydrate (Fig. 1 and 2).

**Fig 1.**
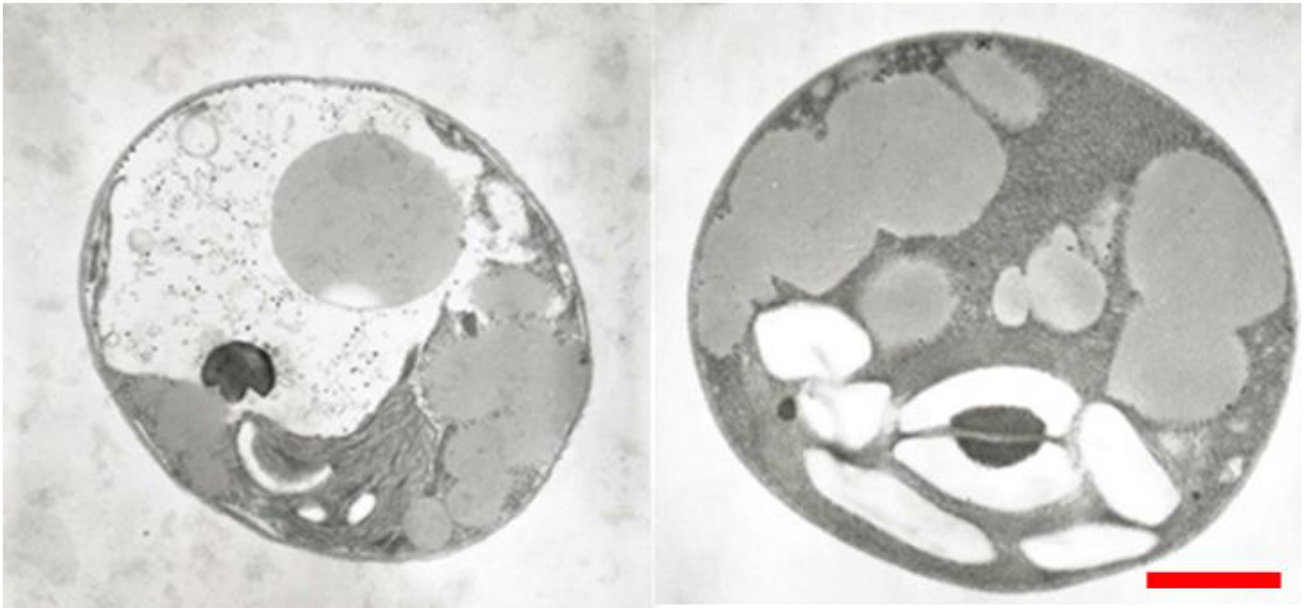
Electron micrographs of *Chlorella* sp. HS2 grown in freshwater (left) and marine (right) growth media at stationary growth phase. Scale bar denotes 1 μm.

**Fig 2.**
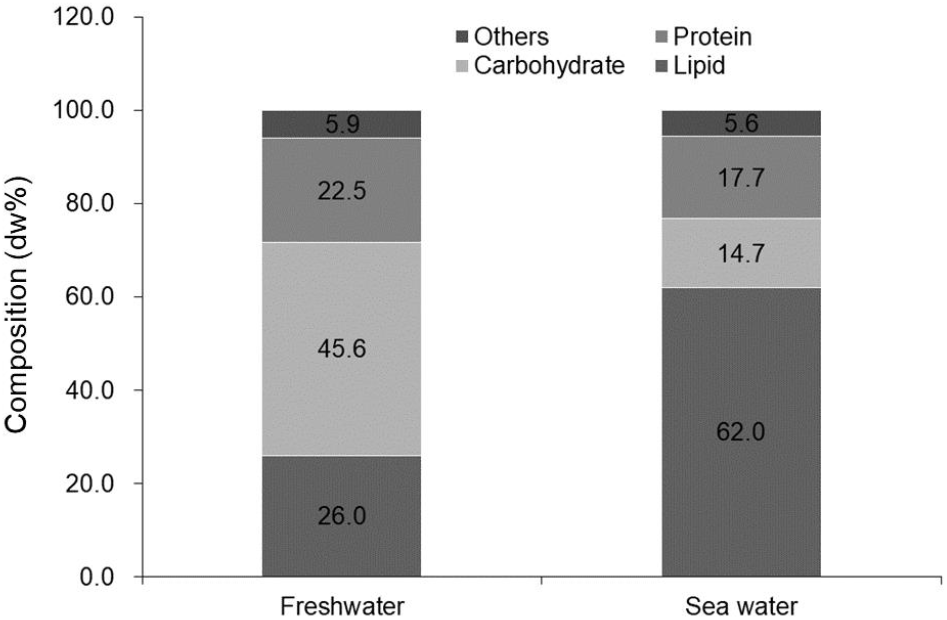
Proximate composition of *Chlorella* sp. HS2 grown in freshwater and marine growth media. Biomass harvested at stationary growth phase (n=3) was used in analysis.

While relatively high amounts of carotenoid pigments (i.e., β-carotene and lutein) under high salinity stress observed in the previous study suggested their possible contribution to the protection of photosynthetic machinery (Talebi, Tabatabaei, Mohtashami, Tohidfar, & Moradi, 2013; Yun et al., 2019), the measures of relative electron transport rate (rETR), the quantum yields of non-photochemical quenching (Y(NPQ)) and non-regulated excess energy dissipation (Y(NO)) using multi-color-PAM indicated that rETR was reduced early during the exponential growth phase under high salinity stress and was recovered at later stationary growth phase. Although differences in Y(NPQ) and Y(NO) were not observed respectively at exponential and stationary phases, a significant difference in Y(NPQ) was observed during stationary phase only at high light intensities and Y(NO) of salt-shocked culture was significantly greater than that of control across all light intensities during exponential growth phase (Fig. 3).

**Fig 3.**
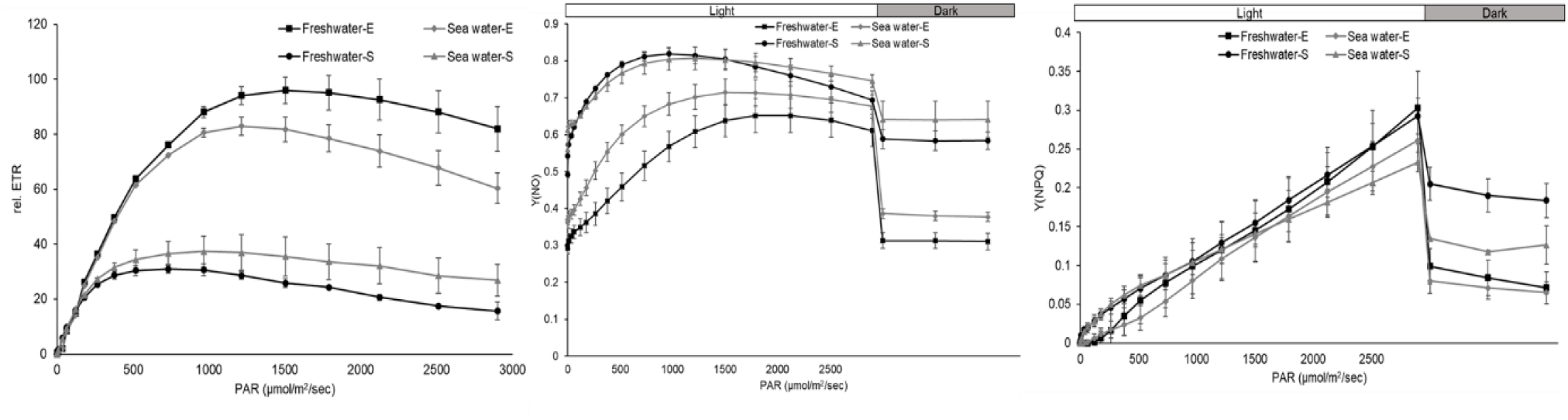
Measurements of parameters related to the photosynthetic activity of *Chlorella* sp. HS2 in freshwater and marine conditions at exponential and stationary growth phases. (a) Relative Electron Transport Rate in PSII. (b) Quantum yield of non-photochemical quenching in PSII. (c) Quantum yield of non-regulated non-photochemical energy loss in PSII. Error bars denote standard error of the mean from triplicate culture samples. E and S respectively denote exponential and stationary growth phases.

### 3.2. Summary of *de novo* assembly

To determine differential transcriptomic regulation of *Chlorella* sp. HS2 under freshwater and marine conditions, RNA-seq was performed using Illumina Hiseq 2000 platform. While *de novo* assembly was performed using Trinity 2.8.5, alignment statistics from Trinity and Bowtie2 2.3.5 mapping results were summarized in Table 1. While 4870 unigenes with 0 count in any of the treatments were filtered out, assessment of assembly quality indicated 89% of complete BUSCOs. Overall, 57640 unigenes were obtained out of 290 million raw reads.

**Table 1.** Statistics of assembled transcriptome of *Chlorella* sp. HS2

| **Statistics of the assembly** |  |
| --- | --- |
| Mean overall alignment rate | 97.14% |
| Unigenes | 57,640 |
| All transcripts | 103,539 |
| Contig N50 | 4,702 |
| Median contig length | 670 |
| Average contig length | 1,948 |
| Percent GC | 63.90% |
| Total assembled bases | 201,680,634 |
| BUSCO results | C:89% [S:20%, D:69%], F:6%, M:4% |

### 3.3. Functional annotation of differentially expressed genes

To elucidate differentially expressed genes (DEGs), read normalization was first performed using SVCD normalization following standard DEGseq2 statistical test; a total of 9117 DEGs were subsequently obtained from 52770 unigenes, and they corresponded to 39469 transcripts. While 3573 DEGs were commonly observed across all conditions, 2334 and 3120 DEGs were distinctively observed at exponential and stationary phases, respectively (Fig. 4). Overall, global observation of transcriptome indicated general transcriptional downregulations under high salinity stress, highlighting substantial metabolic constraints and subsequent biochemical shifts that presumably facilitated the survival of algal cells under high salinity stress. It should be also noted that a substantial difference in terms of the overall DEG expression was observed between exponential and stationary growth phases, with more transcriptional shifts towards downregulation during stationary growth phase. Finally, KO annotation of DEGs yielded 2795 DEGs (i.e., 31% of all DEGs) with 1982 unique consensus KOs, and these DEGs represented one third of genes of *Chlorella variabilis* NC64A’s genome (Eckardt, 2010).

**Fig 4.**
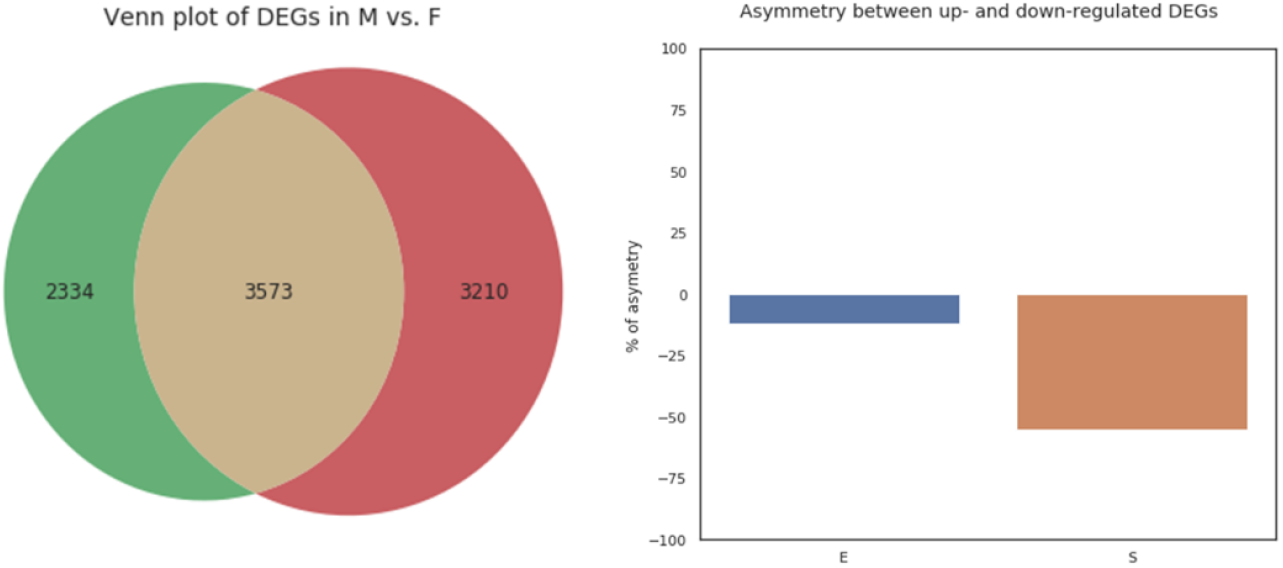
Analysis of DEGs. (A) Venn diagram of the DEGs with an adjusted p-value cutoff of 0.01 in marine (M) and freshwater (F) conditions. (B) Asymmetry between the numbers of up- and downregulated DEGs in exponential (E) and stationary (S) growth phases. Note that negative % asymmetry indicates more DEGs were downregulated generally.

### 3.4. Functional enrichment of differentially expressed genes

Enrichment analysis was performed with the first and second elements of functional hierarchies of KEGG BRITE. While the terms with a p-value lower than 0.05 and a false discovery rate (FDR) lower than 0.25 were considered to be enriched, the results indicated high enrichment of ribosomal proteins (Fig. 5). In addition, papain family of intramolecular chaperones and SNAREs associated with membrane trafficking and heparan sulfate/heparin glycosaminoglycan binding proteins were enriched. Notably, even though FDR values below the cutoff were not observed, many enriched terms with a p-value lower than 0.05 were related to protein processing and membrane trafficking.

**Fig 5.**
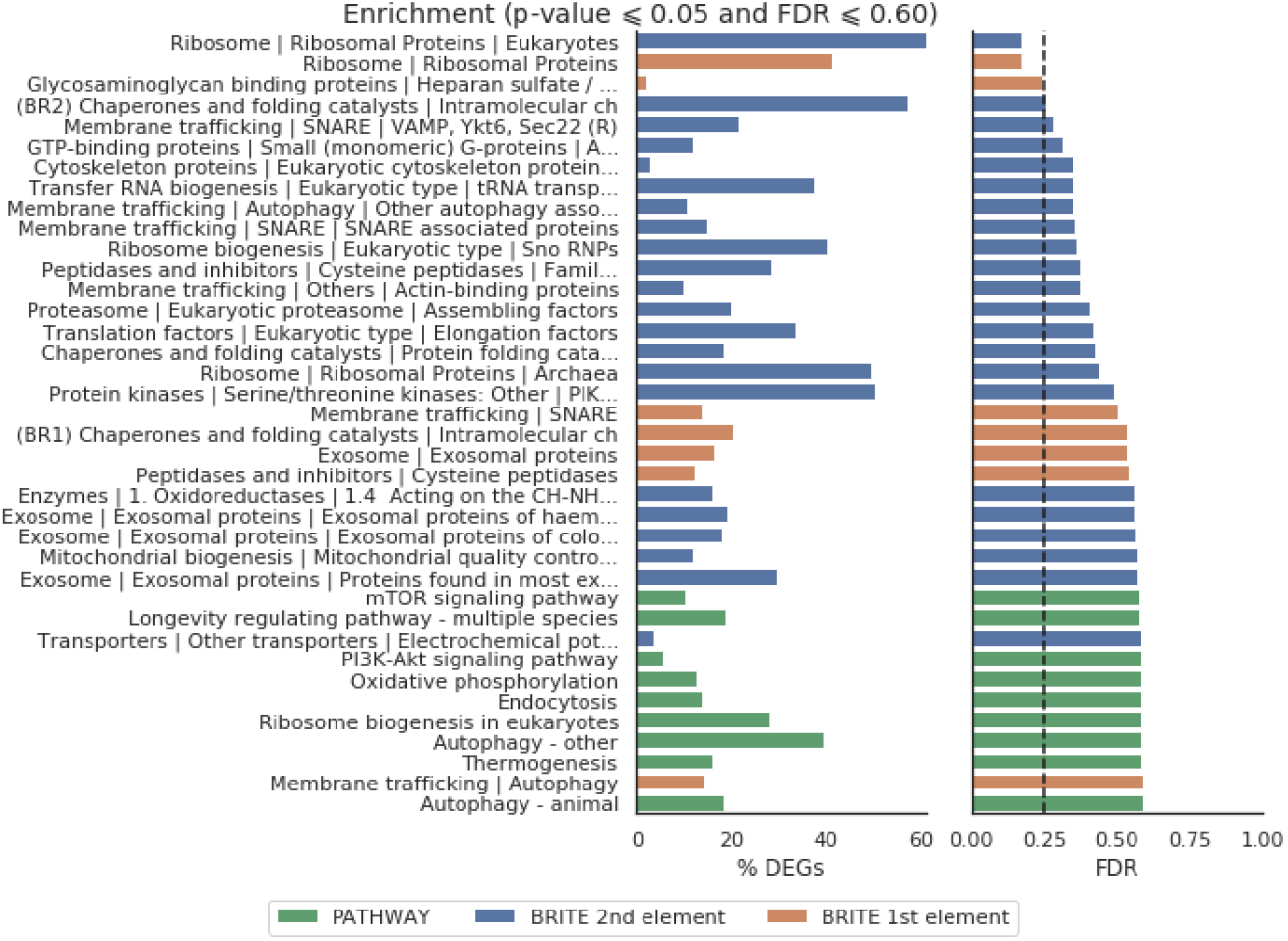
Functional enrichment of DEGs with KEGG pathway and BRITE databases

### 3.5. KEGG pathway analysis

To elucidate metabolic pathways associated with the acclimation of HS2 to high salinity stress, we mapped DEGs to 124 reference pathways in KEGG pathway database. Significantly enriched DEGs were mainly involved in metabolic pathways (cvr01100, 505 unigenes), biosynthesis of secondary metabolites (cvr01110, 238 unigenes), carbon metabolism (cvr01200, 93 unigenes), ribosome (cvr03010, 84 unigenes), biosynthesis of amino acids (cvr01230, 73 unigenes), spliceosome (cvr03040, 53 unigenes), protein processing in endoplasmic reticulum (cvr04141, 44 unigenes), RNA transport (cvr03013, 41 unigenes), glycolysis/gluconeogenesis (cvr00010, 40 unigenes), purine metabolism (cvr00230, 40 unigenes), starch and sucrose metabolism (cvr00500, 38 unigenes), peroxisome (cvr04146, 35 unigenes), oxidative phosphorylation (cvr00190, 33 unigenes), porphyrin and chlorophyll metabolism (cvr00860, 33 unigenes), pyruvate metabolism (cvr00620, 33 unigenes), ubiquitin mediated proteolysis (cvr04120, 32 unigenes), cysteine and methionine metabolism (cvr00270, 29 unigenes), endocytosis (cvr04144, 29 unigenes), aminoacyl-tRNA biosynthesis (cvr00970, 29 unigenes), TCA cycle (cvr00020, 28 unigenes), amino sugar and nucleotide sugar metabolism (cvr00520, 28 unigenes), glyoxylate and dicarboxylate metabolism (cvr00630, 27 unigenes), glycine, serine and threonine metabolism (cvr 00260, 25 unigenes), other types of O-glycan biosynthesis (cvr00514, 25 unigenes), ribosome biogenesis in eukaryote (cvr03008, 25 unigenes), fatty acid metabolism (cvr01212, 24 unigenes), arginine and proline metabolism (cvr00330, 23 unigenes), pyrimidine metabolism (cvr00240, 22 unigenes), photosynthesis (cvr00195, 22 unigenes), sulfur metabolism (cvr00920, 21 unigenes), glutathione metabolism (cvr00480, 20 unigenes), nucleotide excision repair (cvr03420, 20 unigenes), MAPK signaling pathway (cvr04016, 20 unigenes), and RNA degradation (cvr03018, 20 unigenes).

#### 3.5.1. Genes involved in cell cycle and DNA replication

Upon exposure to high salinity stress, the growth of HS2 seemed to be inhibited with an apparent enlargement of cellular biovolume (see 3.1). Correspondingly, most unigenes homologous to genes identified to be involved in cell cycle were downregulated (Table 2). Additionally, DNA replication seemed to be downregulated as well, although Mcm4 of MCM complex (helicase) and DNA polymerase delta subunit 1 [EC: 2.7.7.7] were upregulated (Supplementary Table 2), suggesting the inhibition of DNA replication under high salinity stress. Likewise, most of the unigenes associated with RNA degradation seemed to be downregulated under high salinity stress (Table 2), except CNOT3 (Supplementary Table 1). Furthermore, most genes associated with RNA transport seemed to be downregulated under high salinity stress; and genes associated with aminoacyl-tRNA biosynthesis were downregulated, except glutaminyl-tRNA synthetase [EC: 6.1.1.18] and cysteinyl-tRNA synthetase [EC: 6.1.1.16] (Table 2 and Supplementary Table 1). Although these results generally supported the impairment of both DNA and RNA processing under high salinity stress, it should be emphasized that a number of unigenes associated with repair mechanisms (i.e., nucleotide excision repair, base excision repair, mismatch repair) seemed to be upregulated (Supplementary Table 2).

**Table 2.** Up- and down-regulated genes within KEGG pathway at exponential and stationary growth phases.

| **KEGG pathway** | **Total** | **Upregulation (downregulation)**<br>**at exponential phase** | **Upregulation (downregulation)**<br>**at stationary phase** |
| --- | --- | --- | --- |
| ABC transporters | 16 | 5 (11) | 5 (11) |
| Aminoacyl-tRNA biosynthesis | 43 | 5 (38) | 3 (40) |
| Base excision repair | 15 | 7 (8) | 2 (13) |
| Biosynthesis of unsaturated fatty acids | 8 | 2 (6) | 2 (6) |
| Carbon fixation in photosynthetic organisms | 21 | 5 (16) | 7 (14) |
| Carotenoid biosynthesis | 9 | 3 (6) | 1 (8) |
| Cell cycle | 26 | 8 (18) | 2 (24) |
| Citrate cycle (TCA cycle) | 37 | 8 (29) | 8 (29) |
| DNA replication | 23 | 8 (15) | 2 (21) |
| Fatty acid biosynthesis | 15 | 5 (10) | 4 (11) |
| Fatty acid degradation | 16 | 4 (12) | 1 (15) |
| Glycerolipid metabolism | 13 | 9 (4) | 2 (11) |
| Glycolysis / Gluconeogenesis | 38 | 13 (25) | 12 (26) |
| MAPK signaling pathway | 25 | 4 (21) | 0 (25) |
| Mismatch repair | 11 | 5 (6) | 0 (11) |
| Nucleotide excision repair | 25 | 13 (12) | 4 (21) |
| Photosynthesis | 17 | 1 (16) | 8 (9) |
| Photosynthesis - antenna proteins | 12 | 0 (12) | 7 (5) |
| Proteasome | 32 | 14 (18) | 6 (26) |
| Protein export | 16 | 5 (16) | 1 (15) |
| Protein processing in endoplasmic reticulum | 72 | 20 (52) | 0 (72) |
| Ribosome | 123 | 3 (120) | 2 (121) |
| RNA degradation | 35 | 14 (21) | 5 (30) |
| RNA transport | 62 | 6 (56) | 2 (60) |

#### 3.5.2. Genes involved in protein processing, MAPK signaling pathway, and ABC transporters

While salinity stress is known to inhibit the processing and function of protein, the results indicated the downregulation of enzymes associate with protein processing in endoplasmic reticulum, except mannosyl-oligosaccharide alpha-1,3-glucosidase [EC:3.2.1.207] (GIcII), protein disulfide-isomerase A6 [EC: 5.3.4.1], and protein transport protein SEC24 (Table 2 and Supplementary Table 2). Moreover, most of the ribosomal proteins were downregulated under high salinity stress: of 84 unigenes enriched on KEGG mapper’s ribosome pathway, only S9, S16, and S26e of ribosomal proteins seemed to be upregulated at early exponential growth phase. In addition, while mitogen activated protein kinase (MAPK) signaling cascades are widely recognized for their role in stress response and signal transduction in eukaryotes (Yang, Suh, Kang, Lee, & Chang, 2018), most of the genes associated with MAPK signaling pathway seemed to be downregulated, except P-type Cu^+^ transporter (RAN1) [EC: 7.2.2.8]. Although enriched unigenes indicated that all of the genes associated with protein export were also downregulated under high salinity stress, 3 protein subunits associated with the PA700 (base) of proteasome seemed to be upregulated along with an ABCB subfamily of ABC transporters (i.e., ATM) under high salinity stress (Supplementary Table 1).

#### 3.5.3. Genes involved in photosynthesis and Calvin cycle

There was a clear trend that all of the genes associated with PSII and PSI were downregulated from exponential phase under high salinity stress, corroborating with the measure of PSII activity that indicated a significant reduction in rETR during early growth phase. It should be, however, noted that these genes seemed to be less-downregulated or reverse its downregulation at later stationary growth phase (Table 2 and Supplementary Table 1). Similarly, all of the proteins associated with light harvesting complex (LHC) of HS2 seemed to be downregulated initially under high salinity stress at transcriptional level, whereas Lhcb2 and Lhcb4 were upregulated at the later growth phase. While these results suggested an early compromise in photosynthesis, it should be pointed out that most of the enriched genes involved in carbon fixation via Calvin cycle were downregulated as well (Table 2 and Supplementary Table 1). However, the upregulation of malate dehydrogenase [EC: 1.1.1.40] and alanine transaminase [EC: 2.6.1.2] was observed under high salinity stress and no differential expression in RUBISCO [EC: 4.1.1.39] was observed.

#### 3.5.4. Genes associated with glycolysis and TCA cycle

High salinity stress seemed to induce the upregulation of important genes associated with the conversion of glucose to acetyl-CoA (Table 2 and Supplementary Table 1). In particular, pyruvate dehydrogenase E1 component alpha subunit [EC: 1.2.4.1], which involves in the first step of converting pyruvate to acetyl-CoA was upregulated along with pyruvate decarboxylase [EC: 4.1.1.1]. Moreover, phosphoglucomutase [EC: 5.4.2.2], the enzyme involves in the first step of glycolysis, was upregulated. On the contrary, the results clearly indicated the downregulation of TCA cycle under high salinity stress: most unigenes corresponded to the known genes on TCA cycle were downregulated, suggesting the inhibition of cellular respiration (Table 2 and Supplementary Table 1).

#### 3.5.5. Genes associated with fatty acid and TAG accumulation

Although the genes involved in the synthesis of fatty acids at upstream were downregulated, fatty acyl-ACP thioesterase A [EC: 3.1.3.14] and acyl-desaturase [EC: 1.14.19.2] were upregulated. Provided that the combined amount of C16:1, C18:0, and C18:1 was increased under high salinity stress (Yun et al., 2019), it is especially notable that these two upregulated genes are directly associated with the synthesis of these fatty acids. Moreover, while the genes enriched on KEGG mapper indicated that fatty acid elongation and the biosynthesis of unsaturated fatty acids were not upregulated, survey of fatty acid degradation pathway indicated the inhibition of fatty acid degradation under high salinity stress, suggesting a decrease in fatty acid turnover rate under high salinity stress (Table 2 and Supplementary Table 1).

As the upregulation of lipid synthetic pathway in marine medium was postulated based on the increased lipid content in harvested biomass (see 3.1), the results also identified that genes essential for the synthesis of triacylglycerol (TAG) were upregulated: both phosphatidate phosphate [EC: 3.1.3.4] and diacylglycerol O-acyltransferase 2 [EC: 2.3.1.20] that respectively involve in the conversion of 1,2,-Diacyl-sn-glycerol 3-phosphate to 1,2,-Diacyl-sn-glycerol and in the generation of TAG from 1,2,-Diacyl-sn-glycerol were substantially upregulated under high salinity stress from early growth phase.

#### 3.5.6. Genes associated with carotenoid synthesis

Of 5 unigenes enriched on KEGG mapper’s carotenoid biosynthesis pathway, all of the genes were downregulated, including a gene involved in the conversion of alpha-carotene to lutein (i.e., carotenoid epsilon hydroxylase [EC: 1.14.14.158]) (Supplementary Table 1). In addition, two genes associated with the conversion of phytoene to lycopene, an important intermediate for the synthesis of other carotenoids, were downregulated (i.e., prolycopene isomerase [EC: 5.2.1.13] and zeta-carotene desaturase [EC: 1.3.5.6]) (Supplementary Table 1). Interestingly, both relative and absolute amounts of lutein were increased under high salinity stress (Yun et al., 2019); these results suggest the provision of far-upstream precursors could have played an important role in lutein synthesis.

## 4. Discussion

While high salinity stress strongly influences the viability and biochemical composition of algal crops and thus the economic feasibility of entire algal biorefinery (Kakarla et al., 2018; Laurens et al., 2017; Oh et al., 2019), this study was set out to elucidate transcriptional responses that give rise to the salt tolerance of highly-productive *Chlorella* sp. HS2. Given that genetic engineering approach has been extensively explored with an aim of obtaining robust algal crops, the results clearly indicated that halotolerant HS2 undergoes systematic acclimation responses against high salinity stress, identifying potential target pathways of interest for further genetic modifications or process optimization efforts (Ajjawi et al., 2017; Oh et al., 2019; Qiao, Wasylenko, Zhou, Xu, & Stephanopoulos, 2017). Of these acclimation responses, our results particularly identified that the conversion of stored carbohydrates into algal lipids played a significant role in resolving excess oxidative stress induced under high salinity stress.

These results contradict the overflow hypothesis (OH), which postulates that the overflow of photo-synthase is the main driver of lipid and/or starch accumulation and is frequently discussed to explain increased carbon storage in photosynthetic microbes under stress conditions (Juergens, Disbrow, & Shachar-Hill, 2016; Neijssel & Tempest, 1975; Tan & Lee, 2016): indeed, substantial compromise in PSII activities at both transcriptional and phenotypic levels at early growth phase and the upregulation of enzymes associated with the accumulation of lipid throughout entire growth stages clearly suggest that the overflow of organic photo-synthase *ab initio* was not the main driver of high lipid accumulation under salinity stress. Rather, our results identified the pulling of the acetyl-CoA precursor generated from glycolysis towards lipid synthesis as a major driver of lipid accumulation; accordingly, the redirection of storage carbon likely resulted in the seeming conversion of carbohydrate to lipid as observed in the proximate analysis of harvested biomass. In addition, KEGG pathway analysis of carotenoid synthesis pathway and TCA cycle suggested that these competing pathways for the “pulling” of acetyl-CoA precursor were downregulated, thereby positively contributed to the redirection of acetyl-CoA towards glycerolipid synthesis (Fig. 6).

**Fig 6.**
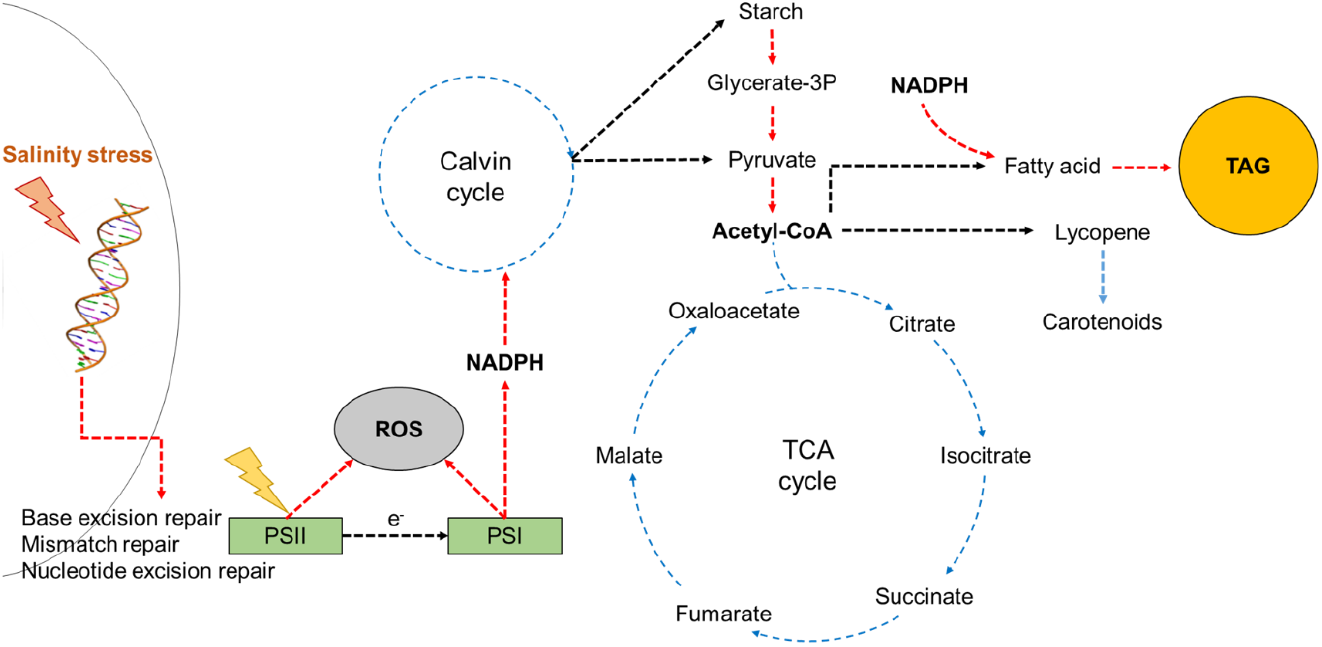
Simplified scheme of carbon and energy flows in *Chlorella* sp. HS2 for putative early responses against high salinity stress. Red and blue dashed arrows respectively indicate upregulation and downregulation of a given conversion or response based on transcriptome or phenotypic analyses. Glycerate-3p Glycerate-3-phosphate; NADPH Nicotinamide adenine dinucleotide phosphate; ROS Reactive Oxygen Species; TAG Triacylglycerol

While Y(NO) of PSII represents non-regulated losses of excitation energy and thus indirectly indicate the relative amount of reactive oxygen species (ROS) (Heinz Waltz GmbH, 2012; Klughammer & Schreiber, 2008), the results suggested strong reduction of PSII accepters and photodamage via formation of ROS during early growth phase under high salinity stress, which seemed to be subsequently resolved at stationary phase. Notably, the downregulation of PsbS, the key protein subunit of PSII regulating conformational changes within PSII light harvesting system that result in energy-dependent quenching (qE), was observed (Kiss, Ruban, & Horton, 2008; Müller, Li, & Niyogi, 2001). However, given that non-photochemical quenching (NPQ) of HS2 under salinity stress was not significantly compromised across both growth phases, it is possible that photoinhibitory quenching (qI) or state transition quenching (qT) were promoted under high salinity stress to compensate for the putative loss of qE in PSII (Zhao et al., 2017). In addition, while high salinity stress was previously known to inhibit the recovery of PSII by damaging D1 protein (Murata, Takahashi, Nishiyama, & Allakhverdiev, 2007), the transcriptome analysis indicated no differential expression of D1 protein of HS2 under high salinity stress. Regardless, overall downregulation of protein processing, including proteasome, under high salinity stress hints at a decrease in D1 protein turnover in PSII (Andersson & Aro, 2001; Erdmann & Hagemann, 2001), which likely contributed to the formation of reactive oxygen species (ROS) and the decreased photosynthetic efficiency at early growth phase (Zhang, Paakkarinen, van Wijk, & Aro, 2000).

In addition to acetyl-CoA precursors, the synthesis of glycerolipid necessitates NADPH as a cofactor (Tan & Lee, 2016). Being an electron donor, NADPH is synthesized along with ATP during the light reaction of photosynthesis, and has been acknowledged for its role as an oxidative stress mediator (Valderrama et al., 2006). Given that KEGG pathway analysis suggested the downregulation of Calvin cycle under high salinity stress, the excess NADPH not utilized in carbon fixation seemed to be directed to the high accumulation of fatty acid and/or glycerolipid, which in turn possibly played an important role in remediating excess oxidative stress at PSII. It should be, however, pointed that common cellular responses under high salinity stress involves the upregulation of anti-oxidative enzymes, including catalase, superoxide dismutase (SOD), and glutathione reductase (GR) as well as DNA repair mechanisms and ABC transporters (Fu, Wang, Yin, Du, & Kan, 2014; Huang, Fulda, Hagemann, & Norling, 2006; Valderrama et al., 2006). Although substantial upregulation of anti-oxidative enzymes was not observed in the results of transcriptome analysis, the degree to which each mitigation response contributes to the overall acclimation of HS2 under high salinity stress across different growth stages remains to be elucidated. Importantly, the upregulation of P-type Cu^+^ transporter (RAN1) on MAPK signaling pathway was observed in HS2 – the activity of RAN1 was determined to be positively correlated with plant cold resistance and the overexpression of RAN1 was further reported to increase abiotic stress tolerance in *Arabidopsis thaliana* (Xu & Cai, 2014; Xu, Zang, Chen, & Cai, 2016; Yang et al., 2018). Furthermore, the increased relative proportion of saturated and mono-saturated fatty acids in HS2 under high salinity stress following the upregulation of enzymes involved in the synthesis of palmitoleate (C16:1), stearate (C18:0), and oleate (C18:1n9c) (Guo, Liu, & Barkla, 2019). Hence, the putative remediation of oxidative stress under growth-inhibiting high salinity condition seems to concurrently involve signal transduction and a shift in membrane fluidity (Guo et al., 2019), in addition to directing excess precursor and co-factor towards lipid synthesis.

While the orchestration of each of elucidated responses likely conferred the relatively high salt tolerance of HS2, lack of some of common algal responses under high salinity stress could offer potential targets in addition to these identified responses when aiming to further enhance the robustness of HS2 as an industrial algal crop. First, violaxanthin deepoxidase (VDE) and zeaxanthin epoxidase (ZEP) that respectively involve in the synthesis of zeaxanthin and violaxanthin were not differentially expressed in HS2 under high salinity stress. Zeaxanthin, however, is known to be associated with several types of photoprotection events of the PSII reaction center (Dall’Osto et al., 2012); therefore, VDE upregulation has been acknowledged as one of common algal responses under high oxidative stress (Z. Li et al., 2016). Given that the relative amount of carotenoid pigments in HS2 was increased under high salinity stress (Yun et al., 2019), enhancing the content of zeaxanthin by either upregulating VDE or downregulating ZEP may further enhance the halotolerance of HS2. Furthermore, although NPQ was not changed under high salinity stress, the elevation of NPQ has been denoted as one of the common algal responses under stress conditions (Cui, Zhang, & Lin, 2017). It would be, therefore, interesting to modulate the NPQ activity of HS2 as part of an effort to confer a greater halotolerance or induce a higher lipid productivity. As an example of the latter, reducing the expression levels of peripheral light-harvesting antenna proteins in PSII was demonstrated to decrease NPQ of *Chlorella vulgaris*, and thereby improved biomass productivity by funneling more photosynthetic energy towards electron transport chain (Shin, Lee, Jeong, Chang, & Kwon, 2016). A similar approach can be adapted to direct more light energy towards electron transport chain and/or possibly increase available NADPH pool, although cautions should be taken to avoid a possibility of antagonistic interactions between competing metabolic pathways.

## Supporting information

Supplementary Table 1. A list of DEGs associated with metabolic pathways of interest.

Supplementary Figure 1. Growth curves of Chlorella Sp. HS2 in terms of DCW and cell density in freshwater and sea water conditions.

Supplementary Table 3. Differential expression analysis results from SVCD-based workflow.

Supplementary Table 4. Trinotate report and lists of aligned transcripts, consensus KOs and aggregated genes by KOs.

Supplementary Table 5. Lists of (a) ambiguous KOs and (b) resolved KOs.

Supplementary File 1. Input file for KEGG mapper search & color.

Supplementary Table 6. Enrichment results by GSEAPY for KEGG Pathway and KEGG BRITE 1st and 2nd elements datasets.

Supplementary Table 2. Raw counts per gene generated by RSEM.

## Acknowledgements

We would like to thank Dr. Carlos P. Roca for his thoughtful comments on SVCD normalization; Sujin Lee and Dr. Saehae Choi for RNA extraction; and Drs. HyunSeok Shin and Byung-Kwan Cho for their suggestions on RNA-seq analysis.

## Notes

**Funding** This work was supported by the Advanced Biomass R&D Center (ABC) of the Global Frontier Program funded by the Ministry of Science and ICT of the Republic of Korea (2016922286), grants from Marine Biotechnology Program and Collaborative Genome Program funded by the Ministry of Oceans and Fisheries of the Republic of Korea (20150184, 20180430), and a grant from KRIBB Research Initiative Program (www.kribb.re.kr).

## References

Ajjawi, I., Verruto, J., Aqui, M., Soriaga, L. B., Coppersmith, J., Kwok, K., … Xu, W. (2017). Lipid production in *Nannochloropsis gaditana* is doubled by decreasing expression of a single transcriptional regulator. Nature biotechnology, 35(7), 647.

Andersson, B., & Aro, E.-M. (2001). Photodamage and D1 protein turnover in photosystem II. In Regulation of Photosynthesis (eds E-M. Aro & B. Andersson), pp. 377–393. Kluwer Academic Publishers, Dordrecht.

Bligh, E. G., & Dyer, W. J. (1959). A rapid method of total lipid extraction and purification. Canadian journal of biochemistry and physiology, 37(8), 911–917.

Buchfink, B., Xie, C., & Huson, D. H. (2015). Fast and sensitive protein alignment using DIAMOND. Nature methods, 12(1), 59.

Camacho, C., Coulouris, G., Avagyan, V., Ma, N., Papadopoulos, J., Bealer, K., & Madden, T. L. (2009). BLAST+: Architecture and applications. BMC bioinformatics, 10(1), 421.

Church, J., Hwang, J.-H., Kim, K.-T., McLean, R., Oh, Y.-K., Nam, B., … Lee, W. H. (2017). Effect of salt type and concentration on the growth and lipid content of *Chlorella vulgaris* in synthetic saline wastewater for biofuel production. Bioresource technology, 243, 147–153.

Cui, Y., Zhang, H., & Lin, S. (2017). Enhancement of non-photochemical quenching as an adaptive strategy under phosphorus deprivation in the dinoflagellate *Karlodinium veneficum*. Frontiers in microbiology, 8, 404.

Dall’Osto, L., Holt, N. E., Kaligotla, S., Fuciman, M., Cazzaniga, S., Carbonera, D., … Bassi, R. (2012). Zeaxanthin protects plant photosynthesis by modulating chlorophyll triplet yield in specific light-harvesting antenna subunits. Journal of biological chemistry, 287(50), 41820–41834.

Dubois, M., Gilles, K. A., Hamilton, J. K., Rebers, P. t., & Smith, F. (1956). Colorimetric method for determination of sugars and related substances. Analytical chemistry, 28(3), 350–356.

Eckardt, N. A. (2010). The *Chlorella* genome: Big surprises from a small package. The Plant Cell, 22, 2924.

Eddy, S. R. (2011). Accelerated profile HMM searches. PLoS computational biology, 7(10), e1002195.

Erdmann, N., & Hagemann, M. (2001). Salt acclimation of algae and cyanobacteria: A comparison. In Algal Adaptation to Environmental Stresses (eds L.C. Rai & J.P Gaur), pp. 323–361. Springer-Verlag Berlin Heidelberg.

Evans, C., Hardin, J., & Stoebel, D. M. (2017). Selecting between-sample RNA-Seq normalization methods from the perspective of their assumptions. Briefings in bioinformatics, 19(5), 776–792.

Flowers, T., Troke, P., & Yeo, A. (1977). The mechanism of salt tolerance in halophytes. Annual review of plant physiology, 28(1), 89–121.

Fogg, G. (2001). Algal adaptation to stress—some general remarks. In Algal Adaptation to Environmental Stresses (eds L.C. Rai & J.P Gaur) pp. 1–19. Springer-Verlag Berlin Heidelberg.

Fu, X., Wang, D., Yin, X., Du, P., & Kan, B. (2014). Time course transcriptome changes in *Shewanella* algae in response to salt stress. PloS one, 9(5), e96001.

Grabherr, M. G., Haas, B. J., Yassour, M., Levin, J. Z., Thompson, D. A., Amit, I., … Zeng, Q. (2011). Trinity: Reconstructing a full-length transcriptome without a genome from RNA-Seq data. Nature biotechnology, 29(7), 644.

Guo, Q., Liu, L., & Barkla, B. J. (2019). Membrane lipid remodeling in response to salinity. International journal of molecular sciences, 20(17), 4264.

Haas, B. J., Papanicolaou, A., Yassour, M., Grabherr, M., Blood, P. D., Bowden, J., … Lieber, M. (2013). De novo transcript sequence reconstruction from RNA-seq using the Trinity platform for reference generation and analysis. Nature protocols, 8(8), 1494.

Heinz Waltz GmbH. (2012). MULTI-COLOR-PAM Manual.

Hu, Q., Sommerfeld, M., Jarvis, E., Ghirardi, M., Posewitz, M., Seibert, M., & Darzins, A. (2008). Microalgal triacylglycerols as feedstocks for biofuel production: Perspectives and advances. The plant journal, 54(4), 621–639.

Huang, F., Fulda, S., Hagemann, M., & Norling, B. (2006). Proteomic screening of salt-stress-induced changes in plasma membranes of *Synechocystis* sp. strain PCC 6803. Proteomics, 6(3), 910–920.

Illman, A., Scragg, A., & Shales, S. (2000). Increase in *Chlorella* strains calorific values when grown in low nitrogen medium. Enzyme and microbial technology, 27(8), 631–635.

Juergens, M. T., Disbrow, B., & Shachar-Hill, Y. (2016). The relationship of triacylglycerol and starch accumulation to carbon and energy flows during nutrient deprivation in *Chlamydomonas reinhardtii*. Plant physiology, 171(4), 2445–2457.

Kakarla, R., Choi, J.-W., Yun, J.-H., Kim, B.-H., Heo, J., Lee, S., … Kim, H.-S. (2018). Application of high-salinity stress for enhancing the lipid productivity of *Chlorella sorokiniana* HS1 in a two-phase process. Journal of microbiology, 56(1), 56–64.

Kent, M., Welladsen, H. M., Mangott, A., & Li, Y. (2015). Nutritional evaluation of Australian microalgae as potential human health supplements. PloS one, 10(2), e0118985.

Kiss, A. Z., Ruban, A. V., & Horton, P. (2008). The PsbS protein controls the organization of the photosystem II antenna in higher plant thylakoid membranes. Journal of biological chemistry, 283(7), 3972–3978.

Klughammer, C., & Schreiber, U. (2008). Complementary PS II quantum yields calculated from simple fluorescence parameters measured by PAM fluorometry and the Saturation Pulse method. PAM application notes, 1(2), 201–247.

Kriventseva, E. V., Kuznetsov, D., Tegenfeldt, F., Manni, M., Dias, R., Simão, F. A., & Zdobnov, E. M. (2018). OrthoDB v10: Sampling the diversity of animal, plant, fungal, protist, bacterial and viral genomes for evolutionary and functional annotations of orthologs. Nucleic acids research, 47(D1), D807–D811.

Langmead, B., Trapnell, C., Pop, M., & Salzberg, S. (2009). Bowtie: An ultrafast memory-efficient short read aligner. Genome Biol, 10(3), R25.

Laurens, L. M., Markham, J., Templeton, D. W., Christensen, E. D., Van Wychen, S., Vadelius, E. W., … Pienkos, P. T. (2017). Development of algae biorefinery concepts for biofuels and bioproducts; a perspective on process-compatible products and their impact on cost-reduction. Energy & Environmental Science, 10(8), 1716–1738.

Lee, H., Nam, K., Yang, J.-W., Han, J.-I., & Chang, Y. K. (2016). Synergistic interaction between metal ions in the sea salts and the extracellular polymeric substances for efficient microalgal harvesting. Algal research, 14, 79–82.

Li, B., & Dewey, C. N. (2011). RSEM: Accurate transcript quantification from RNA-Seq data with or without a reference genome. BMC bioinformatics, 12(1), 323.

Li, Z., Peers, G., Dent, R. M., Bai, Y., Yang, S. Y., Apel, W., … Niyogi, K. K. (2016). Evolution of an atypical de-epoxidase for photoprotection in the green lineage. Nature plants, 2(10), 16140.

Liu, K.-H., Ding, X.-W., Rao, N., Prabhu, M., Zhang, B., Zhang, Y.-G., … Li, W.-J. (2017). Morphological and transcriptomic analysis reveals the osmoadaptive response of endophytic fungus *Aspergillus montevidensis* ZYD4 to high salt stress. Frontiers in microbiology, 8, 1789.

Lowry, O. H., Rosebrough, N. J., Farr, A. L., & Randall, R. J. (1951). Protein measurement with the Folin phenol reagent. Journal of biological chemistry, 193, 265–275.

McBride, R. C., Lopez, S., Meenach, C., Burnett, M., Lee, P. A., Nohilly, F., & Behnke, C. (2014). Contamination management in low cost open algae ponds for biofuels production. Industrial Biotechnology, 10(3), 221–227.

Mootha, V. K., Lindgren, C. M., Eriksson, K.-F., Subramanian, A., Sihag, S., Lehar, J., … Laurila, E. (2003). PGC-1α-responsive genes involved in oxidative phosphorylation are coordinately downregulated in human diabetes. Nature genetics, 34(3), 267.

Müller, P., Li, X.-P., & Niyogi, K. K. (2001). Non-photochemical quenching. A response to excess light energy. Plant physiology, 125(4), 1558–1566.

Murata, N., Takahashi, S., Nishiyama, Y., & Allakhverdiev, S. I. (2007). Photoinhibition of photosystem II under environmental stress. Biochimica et Biophysica Acta (BBA)-Bioenergetics, 1767(6), 414–421.

Neijssel, O., & Tempest, D. (1975). The regulation of carbohydrate metabolism in *Klebsiella aerogenes* NCTC 418 organisms, growing in chemostat culture. Archives of Microbiology, 106(3), 251–258.

Oh, S. H., Chang, Y. K., & Lee, J. H. (2019). Identification of significant proxy variable for the physiological status affecting salt stress-induced lipid accumulation in *Chlorella sorokiniana* HS1. Biotechnology for Biofuels, 12(1), 242.

Peng, Z., He, S., Gong, W., Sun, J., Pan, Z., Xu, F., … Du, X. (2014). Comprehensive analysis of differentially expressed genes and transcriptional regulation induced by salt stress in two contrasting cotton genotypes. BMC genomics, 15(1), 760.

Qiao, K., Wasylenko, T. M., Zhou, K., Xu, P., & Stephanopoulos, G. (2017). Lipid production in *Yarrowia lipolytica* is maximized by engineering cytosolic redox metabolism. Nature biotechnology, 35(2), 173.

Quinn, J. C., & Davis, R. (2015). The potentials and challenges of algae based biofuels: A review of the techno-economic, life cycle, and resource assessment modeling. Bioresource technology, 184, 444–452.

Rathinasabapathi, B. (2000). Metabolic engineering for stress tolerance: Installing osmoprotectant synthesis pathways. Annals of Botany, 86(4), 709–716.

Roca, C. P., Gomes, S. I., Amorim, M. J., & Scott-Fordsmand, J. J. (2017). Variation-preserving normalization unveils blind spots in gene expression profiling. Scientific reports, 7, 42460.

Shin, W.-S., Lee, B., Jeong, B.-r., Chang, Y. K., & Kwon, J.-H. (2016). Truncated light-harvesting chlorophyll antenna size in *Chlorella vulgaris* improves biomass productivity. Journal of Applied Phycology, 28(6), 3193–3202.

Shin, W.-S., Lee, B., Kang, N. K., Kim, Y.-U., Jeong, W.-J., Kwon, J.-H., … Chang, Y. K. (2017). Complementation of a mutation in CpSRP43 causing partial truncation of light-harvesting chlorophyll antenna in *Chlorella vulgaris*. Scientific reports, 7(1), 17929.

Simão, F. A., Waterhouse, R. M., Ioannidis, P., Kriventseva, E. V., & Zdobnov, E. M. (2015). BUSCO: Assessing genome assembly and annotation completeness with single-copy orthologs. Bioinformatics, 31(19), 3210–3212.

Smith, V. H., Sturm, B. S., Denoyelles, F. J., & Billings, S. A. (2010). The ecology of algal biodiesel production. Trends in ecology & evolution, 25(5), 301–309.

Stephens, E., Ross, I. L., King, Z., Mussgnug, J. H., Kruse, O., Posten, C., … Hankamer, B. (2010). An economic and technical evaluation of microalgal biofuels. Nature biotechnology, 28(2), 126.

Subramanian, A., Tamayo, P., Mootha, V. K., Mukherjee, S., Ebert, B. L., Gillette, M. A., … Lander, E. S. (2005). Gene set enrichment analysis: A knowledge-based approach for interpreting genome-wide expression profiles. Proceedings of the National Academy of Sciences, 102(43), 15545–15550.

Talebi, A. F., Tabatabaei, M., Mohtashami, S. K., Tohidfar, M., & Moradi, F. (2013). Comparative salt stress study on intracellular ion concentration in marine and salt-adapted freshwater strains of microalgae. Notulae Scientia Biologicae, 5(3), 309–315.

Tan, K. W. M., & Lee, Y. K. (2016). The dilemma for lipid productivity in green microalgae: Importance of substrate provision in improving oil yield without sacrificing growth. Biotechnology for Biofuels, 9(1), 255.

Unkefer, C. J., Sayre, R. T., Magnuson, J. K., Anderson, D. B., Baxter, I., Blaby, I. K., … Dale, T. (2017). Review of the algal biology program within the National Alliance for Advanced Biofuels and Bioproducts. Algal research, 22, 187–215.

Valderrama, R., Corpas, F. J., Carreras, A., GÓMEZ-RODRÍGUEZ, M. V., Chaki, M., Pedrajas, J. R., … Barroso, J. B. (2006). The dehydrogenase-mediated recycling of NADPH is a key antioxidant system against salt-induced oxidative stress in olive plants. Plant, Cell & Environment, 29(7), 1449–1459.

Valizadeh Derakhshan, M., Nasernejad, B., Abbaspour-Aghdam, F., & Hamidi, M. (2015). Oil extraction from algae: A comparative approach. Biotechnology and applied biochemistry, 62(3), 375–382.

von Alvensleben, N., Stookey, K., Magnusson, M., & Heimann, K. (2013). Salinity tolerance of *Picochlorum atomus* and the use of salinity for contamination control by the freshwater cyanobacterium *Pseudanabaena limnetica*. PloS one, 8(5), e63569.

Wang, L., Yuan, D., Li, Y., Ma, M., Hu, Q., & Gong, Y. (2016). Contaminating microzooplankton in outdoor microalgal mass culture systems: An ecological viewpoint. Algal research, 20, 258–266.

Xu, P., & Cai, W. (2014). RAN1 is involved in plant cold resistance and development in rice (*Oryza sativa*). Journal of experimental botany, 65(12), 3277–3287.

Xu, P., Zang, A., Chen, H., & Cai, W. (2016). The small G protein AtRAN1 regulates vegetative growth and stress tolerance in *Arabidopsis thaliana*. PloS one, 11(6), e0154787.

Yang, A., Suh, W. I., Kang, N. K., Lee, B., & Chang, Y. K. (2018). MAPK/ERK and JNK pathways regulate lipid synthesis and cell growth of *Chlamydomonas reinhardtii* under osmotic stress, respectively. Scientific reports, 8(1), 13857.

Yun, J.-H., Cho, D.-H., Heo, J., Lee, Y. J., Lee, B., Chang, Y. K., & Kim, H.-S. (2019). Evaluation of the potential of *Chlorella* sp. HS2, an algal isolate from a tidal rock pool, as an industrial algal crop under a wide range of abiotic conditions. Journal of Applied Phycology, 31, 1–14.

Yun, J.-H., Cho, D.-H., Lee, B., Kim, H.-S., & Chang, Y. K. (2018). Application of biosurfactant from *Bacillus subtilis* C9 for controlling cladoceran grazers in algal cultivation systems. Scientific reports, 8(1), 5365.

Yun, J.-H., Cho, D.-H., Lee, S., Heo, J., Tran, Q.-G., Chang, Y. K., & Kim, H.-S. (2018). Hybrid operation of photobioreactor and wastewater-fed open raceway ponds enhances the dominance of target algal species and algal biomass production. Algal research, 29, 319–329.

Yun, J.-H., Smith, V. H., La, H.-J., & Keun Chang, Y. (2016). Towards managing food-web structure and algal crop diversity in industrial-scale algal biomass production. Current Biotechnology, 5(2), 118–129.

Yun, J.-H., Smith, V. H., & Pate, R. C. (2015). Managing nutrients and system operations for biofuel production from freshwater macroalgae. Algal research, 11, 13–21.

Zhang, L., Paakkarinen, V., van Wijk, K. J., & Aro, E.-M. (2000). Biogenesis of the chloroplast-encoded D1 protein: Regulation of translation elongation, insertion, and assembly into photosystem II. The Plant Cell, 12(9), 1769–1781.

Zhang, Y., Jiang, Y., & Liang, W. (2006). Accumulation of soil soluble salt in vegetable greenhouses under heavy application of fertilizers. Agricultural Journal, 1, 123–127.

Zhao, X., Chen, T., Feng, B., Zhang, C., Peng, S., Zhang, X., … Tao, L. (2017). Non-photochemical quenching plays a key role in light acclimation of rice plants differing in leaf color. Frontiers in plant science, 7, 1968.

Zhu, L., Wang, Z., Shu, Q., Takala, J., Hiltunen, E., Feng, P., & Yuan, Z. (2013). Nutrient removal and biodiesel production by integration of freshwater algae cultivation with piggery wastewater treatment. Water research, 47(13), 4294–4302.

