## Supplementary figures and images for "Transcriptomic responses associated with carbon and energy flows under high salinity stress suggest the overflow of acetyl-CoA from glycolysis and NADPH co-factor induces high lipid accumulation and halotolerance in *Chlorella* sp. HS2"

### Supplementary Figure 1. Growth curves of Chlorella Sp. HS2 in terms of DCW and cell density in freshwater and sea water conditions.

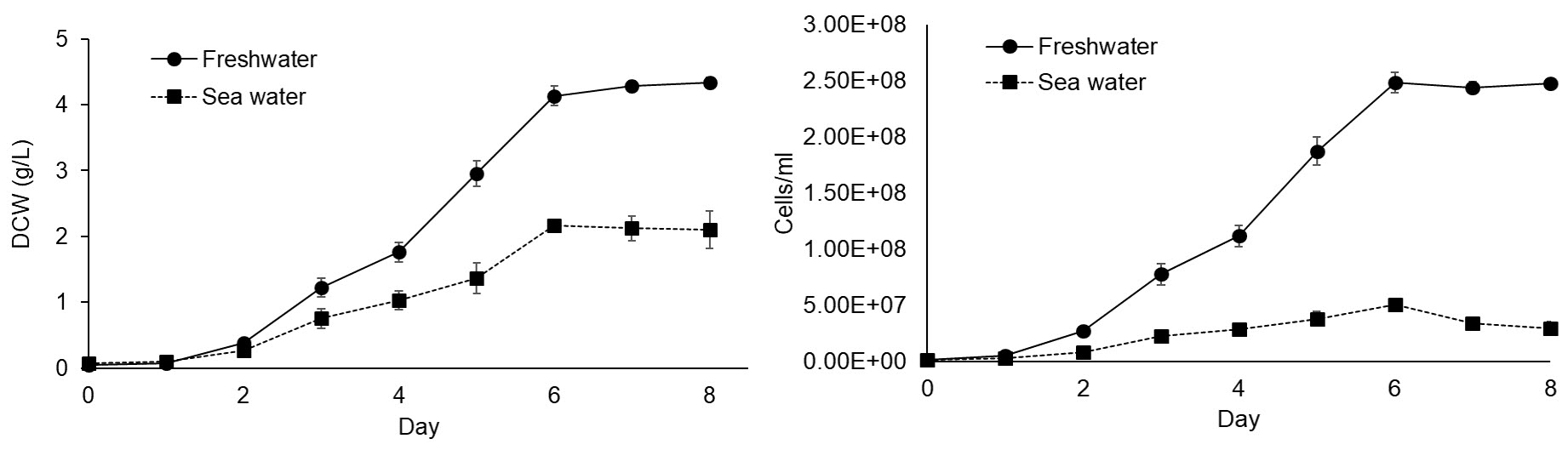
